# Primary restriction of S-RNase cytotoxicity by a stepwise ubiquitination and degradation pathway in *Petunia hybrida*

**DOI:** 10.1101/2020.12.14.422713

**Authors:** Hong Zhao, Yanzhai Song, Junhui Li, Yue Zhang, Huaqiu Huang, Qun Li, Yu’e Zhang, Yongbiao Xue

**Affiliations:** State Key Laboratory of Plant Cell and Chromosome Engineering, Institute of Genetics and Developmental Biology, and The Innovation Academy of Seed Design, Chinese Academy of Sciences, Beijing 100101, China; University of Chinese Academy of Sciences, Beijing 100049, China; Beijing Institute of Genomics, Chinese Academy of Sciences, and National Centre for Bioinformation, Beijing 100101, China; Jiangsu Co-Innovation Center for Modern Production Technology of Grain Crops, Yangzhou University, Yangzhou 225009, China

**Keywords:** ubiquitination, self-incompatibility, S-RNase, SLF, *Petunia hybrida*

## Abstract

In self-incompatible Solanaceous species, the pistil S-RNase acts as cytotoxin to inhibit self-pollination but is polyubiquitinated by the pollen-specific non-self *S*-locus F-box (SLF) proteins and subsequently degraded by the ubiquitin-proteasome system (UPS), allowing cross-pollination. However, it remains unclear how S-RNase is restricted by the UPS. Here, we first show that *Petunia hybrida* (Ph) S_3_-RNase is largely ubiquitinated by K48-linked polyubiquitin chains at three regions, R I, II and III. R I is ubiquitinated in unpollinated, self- and cross-pollinated pistils, indicating its occurrence prior to PhS_3_-RNase uptake into pollen tubes, whereas R II and III are exclusively ubiquitinated in cross-pollinated pistils. Second, removal of R II ubiquitination resulted in significantly reduced seed sets from cross-pollination and that of R I and III in less extents, indicating their increased cytotoxicity. In consistent, the mutated R II of PhS_3_-RNase resulted in marked reduction of its degradation, whereas that of R I and III in less reductions. Taken together, our results demonstrate that PhS_3_-RNase R II functions as a major ubiquitination region for its destruction and R I and III as minor ones, revealing that its cytotoxicity is primarily restricted by a stepwise UPS mechanism for cross-pollination in *P. hybrida*.

**ONE SENTENCE SUMMARY:** Biochemical and transgenic analyses reveal that *Petunia hybrida* S_3_-RNase cytotoxicity is largely restricted by a stepwise ubiquitination and degradation pathway during cross-pollination.

## INTRODUCTION

Self-incompatibility (SI), an inability of a fertile seed plant to produce zygote after self-pollination, represents a reproductive barrier adopted by nearly 40% of flowering plant species to prevent self-fertilization and to promote outcrossing (Nettancourt, 2001). In many species, SI is usually controlled by a single multi-allelic *S*-locus encoding both male and female *S* determinants (Takayama and Isogai, 2005). Their molecular interaction confers the pistil an ability to distinguish between genetically related self- and non-self-pollen. In general, SI can be classified into self- and non-self-recognition systems based on their distinct molecular mechanisms (Fujii et al., 2016). In the self-recognition system of the Papaveraceae and Brassicaceae, self-pollen rejection occurs as a specific interaction between the *S* determinants from the same *S* haplotype. In *Papaver rhoeas*, the female *S*-determinant Prs S (*P*. *rhoeas* stigmatic S) interacts with its cognate Prp S (*P. rhoeas* pollen S) to stimulate a signaling cascade leading to programmed cell death (PCD) of self-pollen (Wilkins et al., 2014). In Brassicaceae, SI response is initiated by the specific interaction of the stigma *S*-locus receptor kinase (SRK) and its cognate pollen coat-localized ligand *S*-locus cysteine-rich protein (SCR/SP11), triggering a phosphorylation-mediated signaling pathway resulting in destruction of factors indispensable for pollen compatibility by the UPS (Samuel et al., 2009). S-RNase-based SI, also termed as Solanaceae-type SI, is a well-studied non-self-recognition system widely present in Solanaceae, Plantaginaceae, Rosaceae and Rutaceae (Fujii et al., 2016; Liang et al., 2020). In this system, the pistil *S* determinant S-RNase serving as cytotoxin can be recognized and ubiquitinated by multiple pollen *S* determinants SLFs forming functional SCF ubiquitin ligases in a collaborative non-self-recognition manner, thus restricting cytotoxicity of non-self S-RNases resulting in cross-pollination (Hua and Kao, 2008; Kubo et al., 2010; Liu et al., 2014; Qiao et al., 2004b; Zhang et al., 2009).

However, it remains largely unclear how S-RNases are specifically regulated in the non-self-recognition system. Currently, two models, the S-RNase degradation model and the S-RNase compartmentalization model, have been proposed to explain how S-RNase cytotoxicity is restricted for cross-pollination (Goldraij et al., 2006; Liu et al., 2014; Qiao et al., 2004b). The degradation model proposed that both self and non-self S-RNases taken up by pollen tubes are mainly localized in the cytosol, where further recognized by SLFs. Entani et al. (2014) showed that SCF^SLF^ complexes can specifically polyubiquitinate non-self S-RNases rather than self ones *in vitro* in *P. hybrida*, providing evidence for S-RNase ubiquitination by cross pollen. Whereas in the self-pollen tubes, binding of self S-RNase and SLF leads to the formation of nonfunctional SCF^SLF^ complex, thus resulting in survival of self S-RNase to inhibit pollen tube growth. Together with the discoveries of SCF^SLF^ complex components such as SLF-interacting SKP1-like 1 (SSK1) and Cullin1 in the species from Solanaceae, Plantaginaceae and Rosaceae (Entani et al., 2014; Huang et al., 2006; Li and Chetelat, 2014; Xu et al., 2013; Zhao et al., 2010), the degradation model appears to function in flowering plants possessing S-RNase-based SI. In *Nicotiana* species, Goldraij et al. (2006) proposed that the majority of self- and non-self S-RNases would be sequestered in vacuole-like structures once imported into pollen tubes and subsequently self recognition between SLFs and a small fraction of S-RNases localized in the cytosol would break the structures, releasing S-RNases in a late-stage of self-pollination, thus triggering the SI response, whereas non-self recognition could stabilize them and maintain S-RNase sequestration. Most previous studies showed that S-RNase degradation instead of compartmentalization acts as the major strategy to restrict S-RNase cytotoxicity (Liu et al., 2014). Nevertheless, little is known about the linkage type of the polyubiquitin chains and the specific residue of S-RNase ubiquitinated by non-self SCF^SLF^ complexes in cross pollen.

To address these questions, in this study, we first established an *in vivo* assay for examining polyubiquitination of PhS_3_-RNase in cross-pollinated pistils and, together with *in vitro* ubiquitination analyses, we found that it is mainly ubiquitinated by K48-linked polyubiquitin chains in three regions named R I, II and III. Among them, R I ubiquitination occurs before PhS_3_-RNase entry into pollen tubes and likely mediated by an unknown E3 ligase, whereas those of R II and III are specifically ubiquitinated by SCF^SLF^. Second, the ubiquitination removal of those three regions had little effect on the physicochemical properties of PhS_3_-RNase, but negatively impacted their functions in cross-pollen tubes. The transgene with a mutated R II led to significant reduction of seed sets from cross-pollination, whereas the mutated R I and III to much less extents in *P. hybrida*, showing that R II ubiquitination of PhS_3_-RNase plays a major role for its destruction and cytotoxicity restriction, whereas R I and III minor roles. Furthermore, the ubiquitination removal of all three regions did not completely inhibit PhS_3_-RNase degradation and cross seed sets, suggesting that the UPS is not the exclusive mechanism to restrict S-RNase cytotoxicity. Taken together, our results demonstrate a stepwise UPS mechanism for primary restriction of S-RNase cytotoxicity during cross-pollination in *P. hybrida*, providing novel mechanistic insight into a dynamic regulation of S-RNases.

## RESULTS

### S-RNase polyubiquitination mainly occurs through K48 linkages at three conserved spatial regions among S-RNases

Previous studies have revealed that S-RNase is ubiquitinated in cross-pollen tubes, but it remains unclear about its ubiquitination linkage type and site. To examine them, we performed an *in vitro* ubiquitination assay and showed that both oligo- and polyubiquitinated PhS_3_-RNase were detected by anti-ubiquitin, -PhS_3_-RNase and -ubiquitin-K48 antibodies compared to *PhS_3_S_3L_* wild-type control, indicating that non-self PhS_3L_SLF1 are capable of forming SCF^SLF^ complex to ubiquitinate PhS_3_-RNase mainly through K48-linked polyubiquitin chains (Fig. 1A). To further detect the ubiquitination site of S-RNase, we used LC-MS/MS and identified six ubiquitinated residues at T102, K103, C118, T153, K154 and K217 of PhS_3_-RNase by wild type pistils cross-pollinated with the transgenic pollen containing the pollen-specific *PhS_3L_SLF1* in *P. hybrida* (Fig. 1B and Supplemental Fig. 1). Furthermore, we found that the ubiquitinated C118, T153, K154 and K217 were exclusively detected in cross-pollinated pistils, suggesting that they are specific for cross-pollination, whereas the ubiquitinated T102 and K103 were detected in both unpollinated and self-pollinated pistils (Supplemental Fig. 2), suggesting that they likely occur before S-RNase uptake into pollen tubes.

To determine the locations of these ubiquitinated amino acid residues in S-RNases, we compared a total of 37 S-RNases from Solanaceae and found that C118 is within conservative (C) 3 region, T102 and K103 adjacent to hypervariable (Hv) b, T153 and K154 between C4 and C5 and K217 at the C terminal region, implying they are located in three largely conserved S-RNase regions (Fig. 1C and Supplemental Fig. 3). Next, we reasoned that the ubiquitination sites should be spatially close to E2. To examine this possibility, we determined the spatial localization of six ubiquitinated residues on the spatial structure of PhS_3_-RNase and found that T102 and K103 are located near the Hvb region on an interface between S-RNase and SLF and termed region (R) I, T153, K154 and K217 in a region close to E2 and termed R II, whereas C118 inside the predicted spatial structure named as R III (Fig. 1D). Taken together, our results demonstrated that S-RNases are ubiquitinated mainly through K48 linkage at three largely conserved spatial regions among S-RNases.

**Figure 1.**
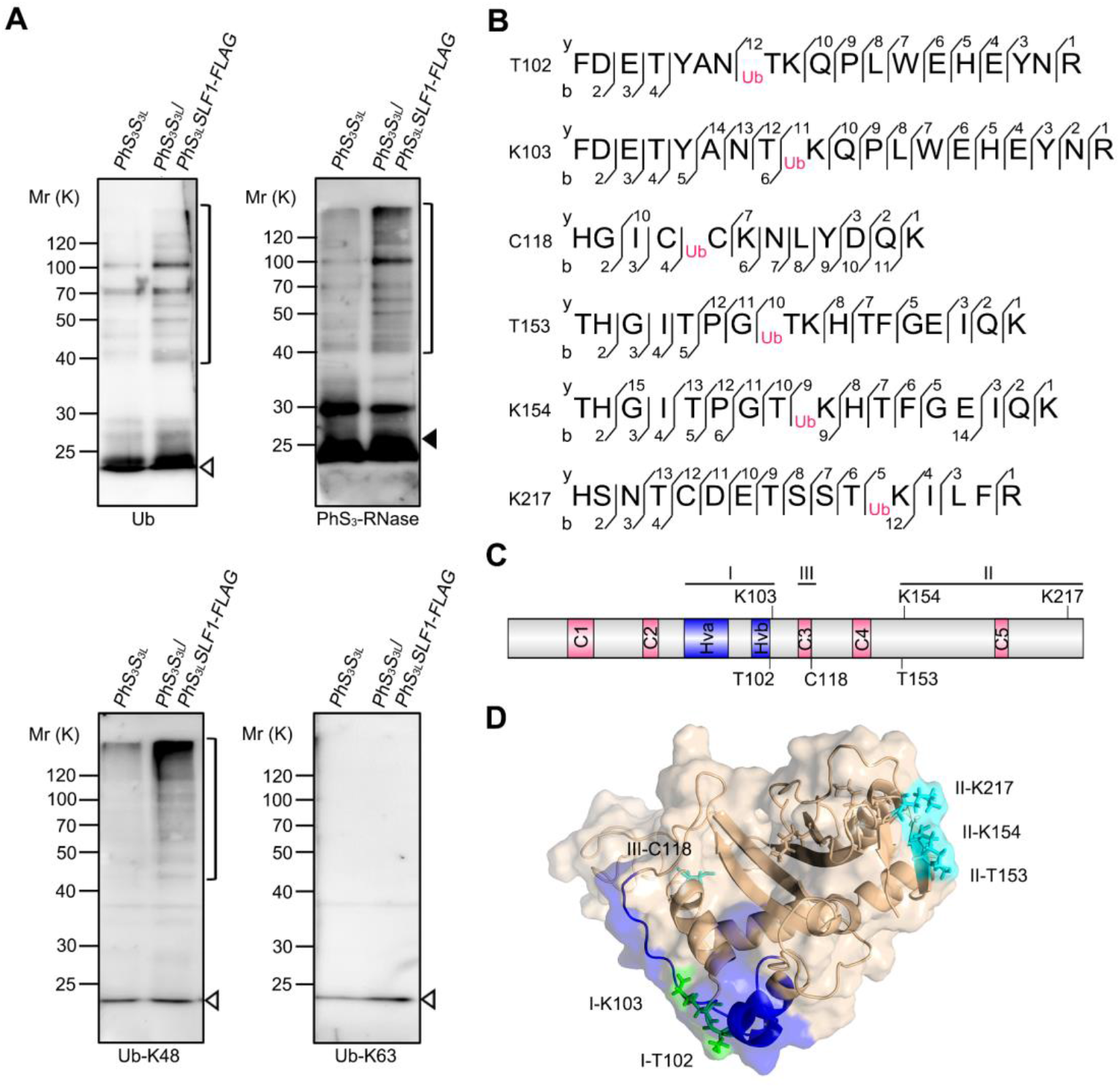
Six amino acid residues of PhS_3_-RNase are ubiquitinated by K48-linked polyubiquitin chains through SCF^Phs_3L_SLF1^. **(A)** Immunoblot detection of *in vitro* ubiquitinated products of PhS_3_-RNase by PhS_3L_SLF1. The pollen genotypes and the transgene are indicated on top and *PhS_3_S_3L_* used as a negative control. Brackets indicate polyubiquitinated PhS_3_-RNases. Open and filled arrowheads indicate ubiquitin and unubiquitinated PhS_3_-RNase monomers, respectively. Antibodies used are indicated in the bottom as Ub, PhS_3_-RNase, Ub-K48 and Ub-K63, respectively. **(B)** Ubiquitination sites of PhS_3_-RNase identified by LC-MS/MS. Ub: the amino acid residue on its right within the peptide sequence of PhS_3_-RNase is ubiquitinated. The b- and y-type product ions are indicated. **(C)** The secondary structural features of PhS_3_-RNase with the locations of the six ubiquitination sites. C1-C5, five conservative regions; Hva and Hvb, hypervariable region a and b. K, C and T: lysine, cysteine and threonine, respectively. **(D)** Spatial locations of the six ubiquitination sites on the 3D structure of PhS_3_-RNase. The dark blue region indicates the Hv regions of PhS_3_-RNase, the green the residues identified by LC-MS/MS in unpollinated, self-pollinated and cross-pollinated pistils, and the cyan the residues identified specifically in cross-pollinated pistils. I, II and III: three regions containing the ubiquitination sites shown in the predicted 3D structure of PhS_3_-RNase.

### Two ubiquitinated amino acids from R I are partially involved in PhS_3_-RNase degradation for cross-pollination

To examine how six ubiquitinated amino acids from three spatial regions mediate the S-RNase ubiquitination, we first designed a mutant construct named MI containing T102A and K103R substitutions incapable of ubiquitination at R I of PhS_3_-RNase and showed that its RNase activity increases with time similar to wild type (Supplemental Fig. 4A), suggesting that MI possesses normal ribonuclease activity. To examine whether the substitutions affect subcellular location of PhS_3_-RNase, we did fractionation experiments and found that MI is predominantly enriched in the S160 fraction derived from the pollen tube cytosol similar to wild type PhS_3_-RNase (S_3_R) (Supplemental Fig. 4B and C). Furthermore, we performed pull-down, split firefly luciferase complementation (SFLC) and bimolecular fluorescence complementation (BiFC) assays and found that MI is capable of interacting with non-self PhS_3L_SLF1 (Supplemental Fig. 4D-F). Nevertheless, we also found that it displayed a weak interaction with self PhS_3_SLF1 (Supplemental Fig. 5), similar to previous studies (Kubo et al., 2010). Consistent with these findings, we found that the structure and electrostatic potentials of MI remain essentially unaltered (Supplemental Fig. 6). Taken together, these results indicated that MI has the enzymatic activity and structure similar to wild type S_3_R.

To examine the *in vivo* function of MI, we transformed *S_3_R* and *MI* driven by the pistil-specific *Chip* promotor into SI *PhS_3_S_3L_* plants, respectively and also transformed their *FLAG*-tagged forms into *PhS_3_S_3L_*. For each construct, we identified at least 24 T_0_ transgenic lines by PCR analysis (Supplemental Fig. 7 and 8) and showed that *MI* is expressed normally in the transgenic lines (Supplemental Fig. 9 and 10A-C). Furthermore, self-pollination assays showed that each construct did not alter the SI phenotypes of the transgenic plants (Supplemental Table 1 and 2). To examine their roles in cross-pollination, we further identified several lines with similar transgene expression levels and found that, compared to about 398 seed set per capsule from *PhS_3_S_3L_* carrying the transgenic *S_3_R* (*S_3_S_3L_/S_3_R*-60) pollinated with cross pollen of *PhS_V_S_V_, S_3_S_3L_/MI* had a reduced seed set of 298 with a reduction of 25% (Supplemental Fig. 10D and E and Supplemental Table 1). In consistent, we also found substantial reduction of seed sets derived from cross-pollination of the *FLAG*-tagged transgenic line *S_3_S_3L_/MI-FLAG*-24 (292 per capsule) with 30% reduction compared to 421 seeds per capsule from *S_3_S_3L_/S_3_R*-*FLAG*-34 (Supplemental Fig. 10D and F and Supplemental Table 2). Taken together, these results suggested that the ubiquitinated R I is involved in cross-pollination.

To verify this role, we assessed the degradation rates of recombinant SUMO-His tagged S_3_R and MI proteins by cell-free degradation assays. As shown in Supplemental Fig. 10G and H, SUMO-His tagged S_3_R degraded rapidly in the samples without MG132 and only 7% was remained after 10-minute treatment. However, the degradation rates were slightly decreased and about 24% remained after 10-minute treatment for SUMO-His-tagged MI protein, showing that MI degradation was partially inhibited and thus resulted in its accumulation in cross-pollen tubes. In addition, degradations of these proteins were significantly delayed by MG132 treatment, indicating the degradation of MI by the UPS pathway similar to wild type.

To confirm the role of the ubiquitinated R I in S-RNase degradation, we further performed ubiquitination assays and found that both polyubiquitinated His-tagged S_3_R and MI proteins were detected by anti-ubiquitin and -PhS_3_-RNase antibodies (Supplemental Fig. 11A), indicating that they both could be ubiquitinated by non-self SCF^PhS_3L_SLF1^. Nevertheless, the amount of ubiquitinated products of MI was reduced to 60% of S_3_R (Supplemental Fig. 11B), suggesting that the ubiquitinated residues located in R I are partially responsible for the ubiquitination of PhS_3_-RNase. Taken together, these results revealed that two ubiquitinated amino acids from R I are partially involved in PhS_3_-RNase degradation for cross-pollination.

### The R II from PhS_3_-RNase serves as a major region for its ubiquitination and degradation in cross-pollen tubes

To examine the function of three ubiquitinated amino acids from R II, we followed a similar strategy to that used for R I by creating MII with T153A, K154R and K217R substitutions of PhS_3_-RNase and found that its RNase activity increases with time similar to wild type (Fig. 2A), suggesting that it possesses normal ribonuclease activity. Second, we found that MII is also predominantly located in the pollen tube cytosol (Fig. 2B), capable of interacting with non-self PhS_3L_SLF1 (Fig. 2C-E), also with a weak interaction with self PhS_3_SLF1 (Supplemental Fig. 5) and had unaltered structure and electrostatic potentials (Supplemental Fig. 6). Furthermore, *MII* and its *FLAG*-tagged transgenes were expressed normally in SI *PhS_3_S_3L_* plants and the transgenic lines also maintained SI phenotype (Fig. 2F-H and Supplemental Fig. 7-9 and Supplemental Table 1 and 2). Compared to *MI* transgenic lines, a significant difference observed for *S_3_S_3L_/MII* was the seed sets derived from pollination with cross pollen of *S_V_S_V_* (ca. 75 seed sets per capsule), with a significant reduction of 81% compared with *S_3_S_3L_*/*S_3_R*-60 (398 seed sets per capsule) (Fig. 2I and J and Supplemental Table 1). In consistent, compared to 421 seeds per capsule from *S_3_S_3L_/S_3_R-FLAG-34*, about 113 seeds were set for the *FLAG*-tagged transgenic lines with a significant 73% reduction (Fig. 2I and K and Supplemental Table 2). We further found that the *MII-FLAG* transgene leads to much less seed set per capsule compared with *MI-FLAG* when their protein levels are similar (Fig. 2H and K), indicating that the ubiquitinated R II plays a major role for cross-pollination. Furthermore, cell-free degradation assay showed that the degradation of MII protein mainly through the UPS pathway is severely inhibited compared with MI in cross-pollen tubes (Fig. 2L and Supplemental Fig. 10H) and *in vitro* ubiquitination assay showed the ubiquitination amount of MII with SCF^PhS_3L_SLF1^ serving as E3 was significantly reduced to 40% of S_3_R, with a reduction of 20% more compared with MI (Supplemental Fig. 11). Taken together, these results suggested that R II of PhS_3_-RNase acts as a major ubiquitination region for its degradation resulting in cross-pollination.

**Figure 2.**
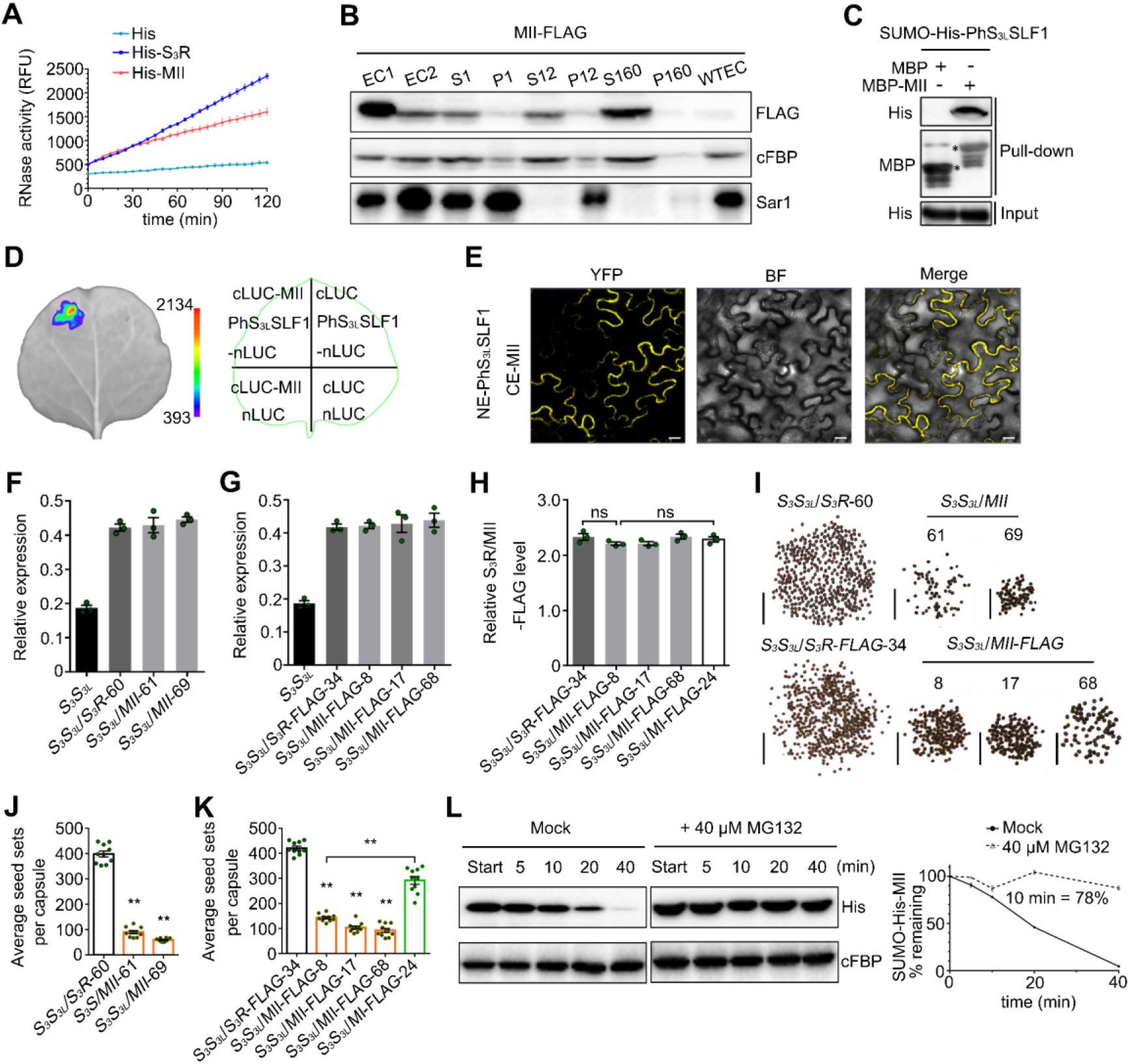
PhS_3_-RNase with mutated R II significantly inhibits cross seed sets. **(A)** RNase activity detection of His-S_3_R and MII expressed by *pCold-TF* vectors. The relative fluorescence unit (RFU) indicating RNase activity during a time course experiment are shown as means ± s.e.m. (n = 3). II: the ubiquitinated region II of PhS_3_-RNase. **(B)** Immunoblot detection of MII-FLAG in subcellular fractions of *in vitro* germinated pollen tubes. EC1, EC2 and WTEC indicate entire cell homogenates of the pistils from the transgenic plants containing *MII*-*FLAG*, the pollen tubes of *PhS_V_S_V_* treated with EC1 and the pistils from wild type *PhS_3_S_3L_*. WTEC was a negative control. S1 and P1, S12 and P12, S160 and P160 indicate supernatant and pellet fractions obtained by centrifugation of EC2 at 1,000 g, 12,000 g and 160,000 g, respectively. cFBP and Sar1 are marker antibodies of cytosol and endoplasmic reticulum (ER), respectively. **(C)** Physical interactions between PhS_3L_SLF1 and MII detected by pull-down assay. Input and pull-down: bait protein SUMO-His-PhS_3L_SLF1 and prey proteins detected by immunoblots, respectively. Asterisks indicate bands of target proteins. **(D)** SFLC assay. The numbers on the left side of the color signal bars represent the values of the fluorescent signal. The injection positions of each component on tobacco leaves are indicated in the contour diagram of leaf margin. **(E)** BiFC assay. NE and CE: transiently expressed N-terminal and C-terminal regions of YFP by *pSPYNE* and *pSPYCE* vectors. YFP, BF and Merge represent the YFP fluorescence, white light and their merged field, respectively. Bars: 20 μm. **(F) and (G)** Transcripts of transgene and native *PhS_3_-RNase* detected by qRT-PCR. The T_0_ transgenic lines are indicated below the horizontal axes. *S_3_S_3L_* is wild type. Data are shown as means ± s.e.m. (n = 3). **(H)** Quantitative analysis of S_3_R- and MII-FLAG proteins. The T_0_ transgenic lines are indicated below the horizontal axes. Data are shown as means ± s.e.m. (n = 3). A student’s *t*-test was used to generate the *p* values. ns, *p* > 0.05; **,*p* < 0.01. **(I)** Reduced seed set per capsule from T0 transgenic lines with mutated R II of PhS_3_-RNase. The transgenic plants containing *S_3_R* or *MII* and *S_3_R-FLAG* or *MII-FLAG* were pollinated with cross pollen of *PhS_v_S_v_*. Numbers below the horizontal lines are T_0_ transgenic line numbers. Bars: 5 mm. **(J) and (K)** Statistical analyses of seed sets per capsule from T_0_ transgenic plants pollinated with cross pollen of *PhS_v_S_v_*. Data are shown as means ± s.e.m. (n ≥ 9). A student’s *t*-test was used to generate the *p* values. **, *p* < 0.01. **(L)** Cell-free degradation of recombinant SUMO-His-MII. Left, immunoblots of the reaction products incubated with or without MG132 (Mock). Start, time point zero in each degradation assay. cFBP antibody was used to detect non-degraded loading control. Right, quantitative analysis of the degradation rates. Data are shown as means ± s.e.m. (n = 3). The remaining amount at 10 min is indicated.

### K154 and K217 from R II act as two major ubiquitination residues for PhS_3_-RNase degradation in cross-pollen tubes

To explore the function of three lysine (K103, K154 and K217) and two threonine (T102 and T153) residues of PhS_3_-RNase in its degradation, we designed two mutant constructs termed MK (K103R, K154R and K217R) and MT (T102A and T153A) which showed similar enzymatic activity, subcellular localization, SLF interactions, structure and electrostatic potentials to wild type S_3_R as well as normal pistil expression and SI phenotype in SI *PhS_3_S_3L_* plants (Fig. 3A-I and Supplemental Fig. 5-9 and Supplemental Table 1 and 2). However, *MK* and *MT* transgenic lines showed differential seed sets of 207 and 356 per capsule after pollination with cross pollen of *S_V_S_V_*, with a significant reduction of 48% and 15%, respectively, compared with *S_3_S_3L_/S_3_R-60* (398 seeds per capsule) (Fig. 3J and Supplemental Fig. 12A and Supplemental Table 1). In consistent, *S_3_S_3L_/MK-FLAG-16* set 113 seeds per capsule with a significant reduction of 73% and 66% more compared with *S_3_S_3L_/S_3_R-FLAG-34* (421 seeds per capsule) and *S_3_S_3L_/MT-FLAG-44* (334 seeds per capsule), respectively (Fig. 3K and Supplemental Fig. 12B and Supplemental Table 2). In addition, we showed that compared with *MT* transgene, *MK* resulted in more seed set reduction similar to *MII* when the transgene expression levels are similar (Fig. 3I and K), suggesting that the identified lysine amino acids especially K154 and K217 play a major role in the ubiquitination and degradation of PhS_3_-RNase. Furthermore, cell-free degradation assays showed that MK degradation by the 26S proteasome had been significantly delayed compared with MT in the absence of MG132 (Fig. 3L and M), and ubiquitination assays indicated that the lysine residues rather than threonine act as the major sites for the PhS_3_-RNase ubiquitination by non-self SCF^PhS_3L_SLF1^ (Fig. 3N). Taken together, our results suggested that K154 and K217 from R II function as two major ubiquitination residues of PhS_3_-RNase for cross-pollination.

**Figure 3.**
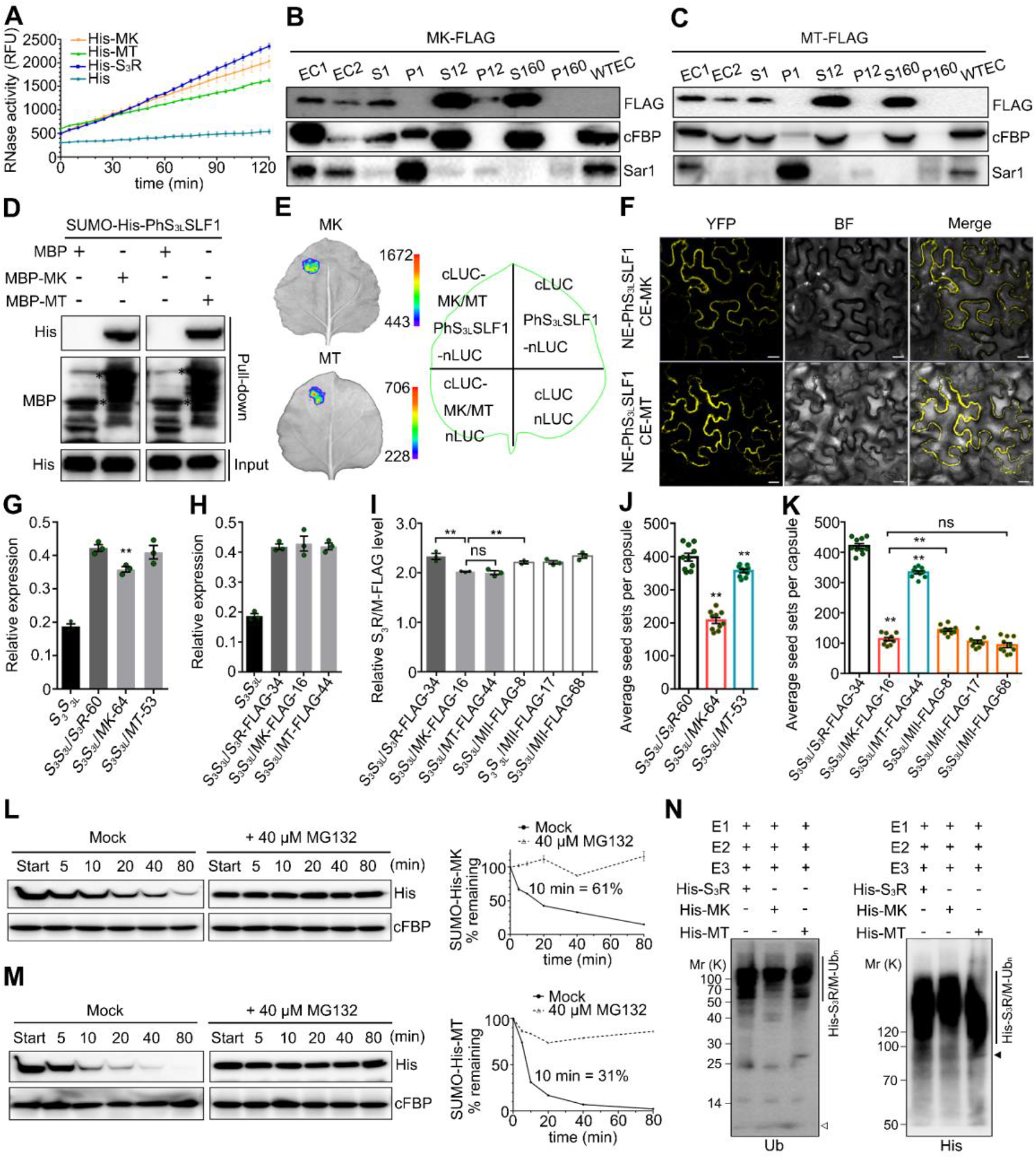
K154 and K217 of the R II of PhS_3_-RNase serve as two major ubiquitination sites for its degradation in cross-pollen tubes. **(A)** RNase activity detection of His-MK and MT expressed by *pCold-TF* vectors. K and T: lysine and threonine within six ubiquitinated residues of PhS_3_-RNase. **(B) and (C)** Immunoblot detection of MK- and MT-FLAG in subcellular fractions of *in vitro* germinated pollen tubes. EC1 and EC2 indicate entire cell homogenates of the pistils from the transgenic plants containing *MK*- **(B)** or *MT*-*FLAG* **(C)**, and the pollen tubes of *PhS_V_S_V_* treated with EC1. **(D)** Physical interactions between PhS_3L_SLF1 and MK and MT detected by pull-down assays. **(E)** SFLC assays. **(F)** BiFC assays. **(G) and (H)** Transcripts of transgene and native *PhS_3_-RNase* detected by qRT-PCR. Data are shown as means ± s.e.m. (n = 3). **,*p* < 0.01. **(I)** Quantitative analysis of S_3_R-, MK- and MT-FLAG proteins. Data are shown as means ± s.e.m. (n = 3). **(J) and (K)** Statistical analyses of seed sets per capsule from T_0_ transgenic plants pollinated with cross pollen of *PhS_v_S_v_*. Data are shown as means ± s.e.m. (n = 10). **(L) and (M)** Immunoblots of recombinant SUMO-His-MK and -MT in the cell-free degradation products incubated with or without MG132 (Mock). **(N)** Immunoblots detection of *in vitro* ubiquitination products of His-S_3_R, -MK and -MT by sCF^PhS_3L_SLF1-FLAG^ (E3) using anti-Ub and -His antibodies. The vertical lines illustrate the ubiquitinated substrates. Open and filled arrowheads indicate ubiquitin and unubiquitinated substrate monomers, respectively. Annotations of this figure are identical to those of Figure 2.

### R III functions as the second major ubiquitination region for PhS_3_-RNase degradation allowing cross-pollination

To investigate the function of the ubiquitination site C118 from the internal R III, we designed MIII (C118A) and found that it also maintains ribonuclease activity, subcellular localization and structure similar to wild type S_3_R (Supplemental Fig. 5, 6 and 13). We further transformed *MIII* and its *FLAG*-tagged form into SI *PhS_3_S_3L_* plants and detected significantly reduced seed sets of about 160 and 261 per capsule from *S_3_S_3L_/MIII*-84 and *S_3_S_3L_/MIII-FLAG*-18 after pollination with cross pollen of *S_V_S_V_*, with a reduction of 59% and 38% compared with *S_3_S_3L_/S_3_R*-60 and *S_3_S_3L_/S_3_R-FLAG-34*, respectively (Supplemental Fig. 7-9 and 14A-G and Supplemental Table 1 and 2). Furthermore, the average seed set per capsule was much less than *S_3_S_3L_/MI-FLAG* when they showed similar transgene expression levels (Supplemental Fig. 14C, F and G), supporting a role of R III in the degradation of PhS_3_-RNase. In addition, we detected a marked accumulation of MIII in cross-pollen tubes compared with S_3_R and MI in the absence of MG132 (Supplemental Fig. 10G, H and 14H) and a significant decreased ubiquitination amount similar to MII (Supplemental Fig. 11). Taken together, these results suggested that the R III acts as a second major ubiquitination region for the degradation of PhS_3_-RNase leading to cross-pollination.

### R I, II and III of PhS_3_-RNase function additively in its degradation for cross-pollination

To examine the function of the three ubiquitination regions together, we made MI/II/III (T102A, K103R, T153A, K154R, K217R and C118A). Similar to wild type PhS_3_-RNase, the mutant form exhibited normal physicochemical properties (Supplemental Fig. 5, 6 and 15) but resulted in 197 and 93 cross seeds per capsule derived from *S_3_S_3L_/MI/II/III-45* and *S_3_S_3L_/MI/II/III-FLAG-49* with pollen of *S_V_S_V_*, respectively, a significant reduction of 50% and 77% similar to the lines containing the mutated R II (Fig. 4A-G and Supplemental Fig. 7-9 and Supplemental Table 1 and 2). Furthermore, the degradation of MI/II/III in cross-pollen tubes was strongly inhibited in 40 min in the absence of MG132 (Fig. 4H and I), indicating a significantly reduced ubiquitination by SCF^PhS_3L_SLF1^ (Fig. 4J). Taken together, these results suggested that the degradation of PhS_3_-RNase is largely dependent on an additive role of its three ubiquitination regions for cross-pollination.

**Figure 4.**
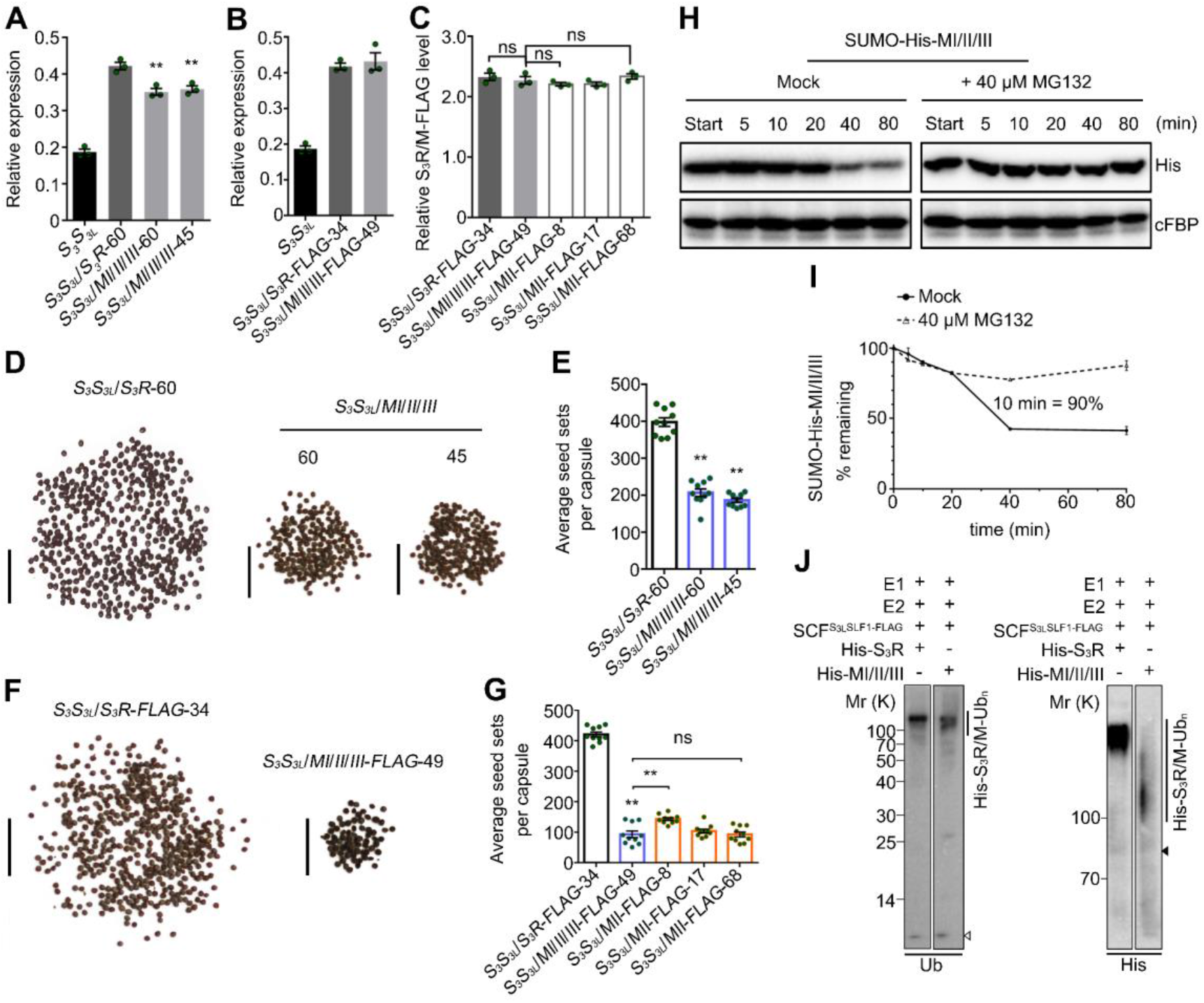
PhS_3_-RNase with mutated R I, II and III regions significantly reduces cross seed sets. **(A) and (B)** Transcripts of transgene and native *PhS_3_-RNase* detected by qRT-PCR. Data are shown as means ± s.e.m. (n = 3). I, II and III, three ubiquitination regions of PhS_3_-RNase. **(C)** Quantitative analysis of S_3_R- and MI/II/III-FLAG proteins. Data are shown as means ± s.e.m. (n = 3). **(D)** Reduced seed set per capsule from cross-pollinated T_0_ transgenic lines containing *MI/II/III.* The transgenic plants containing *MI/II/III* were pollinated with cross pollen of *PhS_v_S_v_*. Numbers below the horizontal lines indicate T_0_ transgenic line numbers. **(E)** Statistical analyses of seed sets per capsule from T_0_ transgenic lines described in **D**. Data are shown as means ± s.e.m. (n = 10). **(F)** Reduced seed set per capsule from cross-pollinated T_0_ transgenic lines containing *MI/II/III-FLAG*. **(G)** Statistical analyses of seed sets per capsule from T_0_ transgenic lines described in **F**. Data are shown as means ± s.e.m. (n = 10). **(H)** Immunoblots of recombinant SUMO-His-MI/II/III in the cell-free degradation products incubated with or without MG132 (Mock). **(I)** Quantitative analyses of immunoblots from **H** and the degradation rate. **(J)** Immunoblot detections of *in vitro* ubiquitinated recombinant His-tagged S_3_R and MI/II/III. Annotations of this figure are identical to those of Figure 2 and 3.

## DISCUSSION

Previous studies have shown that non-self S-RNases are collaboratively recognized by multiple non-self SLFs leading to the formation of canonical SCF^SLF^ complexes for their ubiquitination and subsequent degradation by the 26S proteasome resulting in cross-pollination, but the ubiquitination linkage type and site remain unclear. In this study, we have found that non-self S-RNase is mainly polyubiquitinated through K48 linkages by SCF^SLF^ at three spatial regions (R I, II and III) in *P. hybrida*. Among them, R I ubiquitination appears to occur before S-RNase uptake into pollen tubes with a minor role, if any, in cross-pollen tubes, whereas R II and III act as two major ubiquitination regions for S-RNase degradation. Based on our results, we propose a stepwise UPS model for S-RNases cytotoxicity restriction allowing cross-pollination in *P. hybrida* (Fig. 5). In this model, both self and non-self S-RNases with a small fraction of R I ubiquitinated forms likely mediated by an unknown E3 ligase are taken up into the cytosols of either self-or cross-pollen tubes. Firstly, the R I ubiquitinated forms would make them unable to be recognized by SLFs but degraded by the 26S proteasome. Secondly, other S-RNases could be recognized by SLFs on the basis of ‘like charges repel and unlike charges attract’, and the like electrostatic potentials together with other unknown forces between self S-RNase and its cognate SLF would result in formation of non-functional SCF^SLF^ complexes as demonstrated previously (Li et al., 2017), whereas non-self S-RNase would be attracted by unlike electrostatic potentials and other unknown factors and polyubiquitinated by functional SCF^SLF^ complexes at R II leading to its degradation by the 26S proteasome. Thirdly, the internal R III of non-self S-RNase could be exposed by a conformational change for its further ubiquitination by SLFs and degradation resulting in cross-pollination. Our studies have revealed that the ubiquitination and degradation of non-self S-RNases depend on at least three regions with distinct ubiquitination sites including lysine, threonine and cysteine, reinforcing the notion that the restriction of S-RNase cytotoxicity occurs mainly by the ubiquitination-mediated degradation mechanism (Liu et al., 2014).

**Figure 5.**
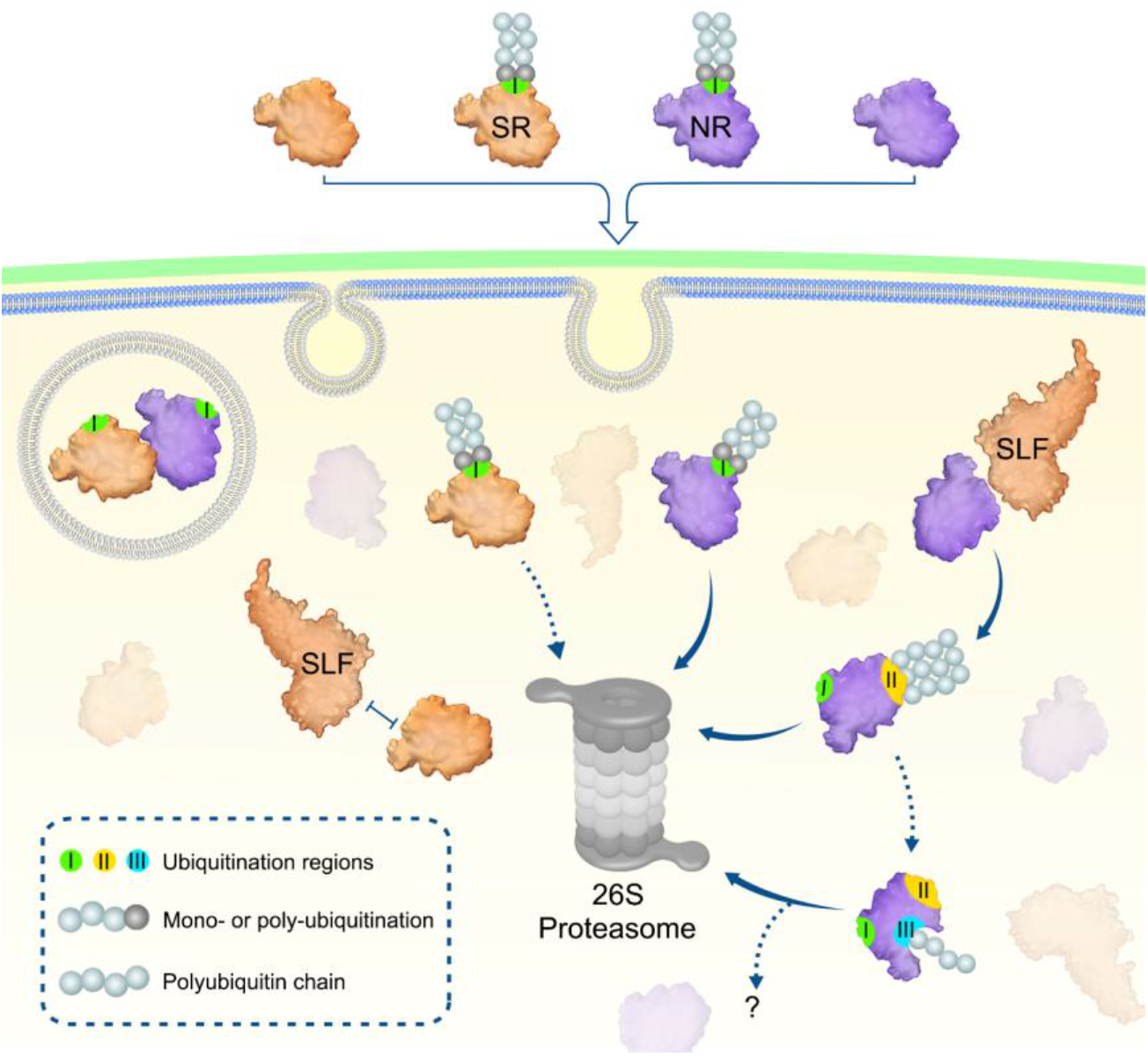
Proposed model for a stepwise ubiquitination and degradation mechanism of S-RNases. SR, NR and their R I ubiquitinated forms can enter the cytoplasm through the pollen tube membrane. The R I ubiquitinated S-RNases could be degraded by the 26S proteasome, whereas their unubiquitinated forms are identified by SLFs. SR and its cognate SLF repel each other and results in an inability of SCF^SLF^ to ubiquitinate SR. In contrast, NR is attracted by non-self SLF for R II ubiquitination by SCF^SLF^ and its subsequent degradation by the 26S proteasome. Subsequently, internal R III could be further exposed for ubiquitination, leading to degradation of NR by the 26S proteasome or other unknown pathway. In addition, S-RNase compartmentalization could occur in the vacuole and contribute to its sequestration. SR and NR: self and non-self S-RNases; I, II, and III: three ubiquitination regions in S-RNases.

Nevertheless, the underlying mechanisms of R I and III ubiquitination remain to be further elucidated. Notably, newly synthesized secretory proteins are constantly scrutinized and destructed by the quality control system such as ER-associated degradation (ERAD) or autophagy to maintain proteostasis once they are misfolded or aggregated (Anelli and Sitia, 2008). The ubiquitination of R I in the unpollinated pistils suggested that it might be resulted from the polyubiquitination of misfolded PhS_3_-RNases. Ubiquitination of closely spaced residues is predicted to be important for polyubiquitin chain assembly (Wang et al., 2012). Here, we found that the identified threonine ubiquitination residues paired with lysine also contribute to the ubiquitination and degradation of non-self S-RNase for cross-pollination, suggesting that they may play a role in building polyubiquitin chains long enough for proteasome recognition (Thrower et al., 2000). In addition, ubiquitin often serves as a critical signal governing the membrane traffic system. Monoubiquitination is sufficient to initiate the internalization of plasma membrane proteins, and K63-linked polyubiquitination is frequently involved in their subsequent sorting and trafficking (Paez Valencia et al., 2016). It is thus possible that R I could also be monoubiquitinated leading to S-RNase entry into pollen tubes by endocytosis, consistent with our results showing that a small fraction of S-RNases are sequestered in microsome fractions. As for the internal R III, it is possible that it would be ubiquitinated unless exposed. Thus, the ubiquitination of R II might lead to the conformational change of non-self S-RNase and exposure of R III, and the subsequent ubiquitination of R III would further block the enzymatic activity of the S-RNase (Sagar et al., 2007). Previous simulations demonstrated that the conserved complementary electrostatic patterns and hydrophobic patches of Rpn10, a recognition subunit of proteasome, and K48-linked tetraubiquitin of the substrates are critical for their interaction (Zhang et al., 2016). Likewise, the ubiquitination of R III might further enhance the electrostatic potentials and hydrophobicity to strengthen the recognition of ubiquitinated S-RNase by the proteasome as well as its degradation.

It remains unclear why the additive action of the three regions did not completely restrict non-self S-RNase cytotoxicity in cross-pollen tubes. We suggest that there might be additional mechanism (s) occurring either after the R III-mediated ubiquitination of non-self S-RNase or during S-RNase uptake into pollen tubes by endocytosis. In *Nicotiana*, S-RNases appeared to be sequestered early in vacuole but released out late to the cytosol in self-pollen tubes (Goldraij et al., 2006). In *P. hybrida*, a small amount of S-RNases are detected in the microvesicles of pollen tubes (Liu et al., 2014). Together, these findings suggest that S-RNase compartmentalization could function during an early phase of S-RNase action. Further investigation into S-RNase uptake mechanism would provide answer to this possibility.

T2 RNases are widespread in every organism except Archaea and involved in a variety of biological processes, including phosphate starvation, viral infection, self-fertilization, tumor growth control and cell death (Bariola et al., 1994; Löffler et al., 1992; Meyers et al., 1999; Ramanauskas and Igić, 2017; Thompson and Parker, 2009). However, our understanding of their functions remains largely incomplete, especially in the case where their roles appear to be independent of their enzymatic activity. In *Saccharomyces cerevisiae*, T2 RNase Rny1 can be released from the vacuole to cleave tRNA and rRNA under superoxygen stress (Thompson and Parker, 2009). Rny1 is indispensable for cell viability, but overexpressed Rny1 can act as a cytotoxin during oxidative stress (MacIntosh et al., 2001; Thompson and Parker, 2009). Moreover, its inactivation strikingly has no effects on cell viability (MacIntosh et al., 2001), but the underlying mechanism remains elusive. In human, RNASET2 is not only implicated to regulate neurodevelopment downstream immune response, but also serve as a tumor suppresser (Henneke et al., 2009), whereas how it contributes to this process in a cleavage-independent manner is poorly defined. In addition, the catalytic-independent function of T2 RNase has also been confirmed for ACTIBIND from *Aspergillus niger* that can bind and destroy the normal actin networks, which is supposed to be conserved in other T2 RNase family members including S-RNase (Roiz et al., 2006). Thus, T2 RNase may act as a molecular signal mediating multiple biological settings, revealing that diverse T2 RNase roles could be derived through neofunctionalization in these lineages.

Distinct from S-RNase-based SI widely present in Solanaceae, Rosaceae, Plantaginaceae and Rutaceae, SI in Brassicaceae and Papaveraceae is well-characterized with a signaling transduction pathway stimulated by the self-recognition between the pistil and pollen *S* determinants. In Brassicaceae, the binding between the pollen-specific SCR/SP11 and its cognate pistil-specific SRK can induce homodimerization and autophosphorylation of SRK, thus triggering the phosphorylation and activation of MLPK to further promote the phosphorylation-mediated recruitment of ARC1 E3 ligase. Consequently, the SI inhibitors or SC factors like EXO70A1 are ubiquitinated and degraded leading to self-pollen rejection (Samuel et al., 2009). The SI-induced signaling in *P. rhoeas* is mainly focused on the initiation of PCD. During the early period after self-pollination, an influx of Ca^2+^ and K^+^ is stimulated, triggering a signaling cascade including phosphorylation and inactivation of Pr-26.1a/b, phosphorylation and activation of p56, F-actin depolymerization and increased ROS and NO (Wilkins et al., 2014). Among them, the disrupted cytoskeleton dynamic serves as a major cause of PCD (Thomas et al., 2006), which is also proposed to occur in self-pollen tubes of *Pyrus bretschneideri* (Chen et al., 2018). Yang et al. (2018) reported an S-RNase-mediated actin disruption in apple (*Malus×domestica*) (Yang et al., 2018). Moreover, self S-RNase can disrupt Ca^2+^ gradient at pollen tube apex through inhibiting phospholipase C (PLC) (Qu et al., 2017). In addition, heat-inactivated S-RNase surprisingly exerts a more sever inhibition of pollen tubes (Gray et al., 1991). These studies suggest that S-RNase could function in a signaling pathway independent of its enzymatic activity. Our results indicated that self S-RNase could be partially ubiquitinated extracellularly to be destroyed during its uptake, but we can not rule out the possibility that its ubiquitination could act as an initial signal for SI response. In addition to ubiquitination, a recent study in *Solanum chacoense* showed that the numbers of carbohydrate chains of S-RNases may influence pollen rejection threshold of S-RNase (Liu et al., 2008). As phosphorylation serves as a critical modification modulating multiple cellular events, it may also be involved in S-RNase activity regulation and the downstream signaling transduction in Solanaceae-type SI. Besides, previous studies have shown that there should exist other factors except electrostatic potentials contribute to the recognition between SLF and S-RNase (Li et al., 2017). Thus, future studies on the structure of SLF bound to S-RNase, other post-translational modifications such as glycosylation and phosphorylation of S-RNase and their relationships with its ubiquitination should shed light on how S-RNase functions and stimulates downstream signaling networks in the pollen tubes.

In sum, our results have revealed a novel stepwise UPS mechanism for S-RNase cytotoxicity restriction resulting in cross-pollination in *P. hybrida*. Our findings also indicate a possible mechanism for dynamic regulation of secreted cytotoxin activities including other T2 ribonuclease members. Further validation of this mechanism using biochemical and cytological approaches is expected to provide additional insights into T2 RNase neofunctionalization.

## METHODS

### Plant materials

Self-incompatible wild-type lines of *PhS_3_S_3_, PhSvSv* and *PhS_3_S_3L_* have been previously described (Robbins et al., 2000; Sims and Ordanic, 2001). The transgenic plants *PhS_3_S_3L_/PhS_3L_SLF1-FLAG* were constructed by transforming *PhS_3_S_3L_* with *pBI101-PhS_3_A-SLF::PhS_3L_SLF1-FLAG. PhS_3_A-SLF* is a native promotor used for *PhSLFs* expression as previously described (Liu et al., 2014; Qiao et al., 2004a).

### Ti plasmid construction and transgenic plant generation

*PhS_3_-RNase* cDNA and its *FLAG*-tagged form were amplified by primers listed in Supplemental Table 3 to introduce *XhoI* and *SacI* restriction sites at their 5’ and 3’ end, respectively. *PhS_3_-RNase* point mutations were generated by PCR (Polymerase chain reaction) using site-directed mutagenesis primers listed in Supplemental Table 3 with *pEASY-PhS_3_-RNase* or *pEASY-PhS_3_-RNase-FLAG* construct as the template. *MI* was generated by mutating T102 and K103 into alanine and arginine, *MII* T153 and K154 into alanine and arginine, and K217 into arginine, *MIII* C118 into alanine, *MK* K103, K154 and K217 into arginine, *MT* T102 and T153 into alanine on *PhS_3_-RNase, MI/II/III* T102, K103 and C118 into alanine, arginine and alanine, respectively on *MII.* The *GUS* gene of *pBI101* was removed using double-digestion, and the pistil-specific promotor *Chip* was ligated into *pBI101* by *KpnI* and *XhoI. PhS_3_-RNase*-(*FLAG*) and its six mutant (m) forms were digested with *Xho*I and *Sac*I and inserted into *pBI101* containing *Chip* promotor to generate *pBI101-Chip::PhS_3_-RNase-(FLAG)* and *pBI101-Chip::PhS_3_-RNase (m)-(FLAG)*. The vectors were separately electroporated into *Agrobacterium tumefaciens* strain LBA4404 and introduced into *PhS_3_S_3L_* using the leaf disk transformation method described previously (Lee et al., 1994; Qiao et al., 2004a).

### Protein structure prediction and electrostatic potential analysis

PhS_3_-RNase protein structure was modeled by the I-TASSER server (http://zhanglab.ccmb.med.umich.edu/I-TASSER/) according to the instructions (Yang et al., 2015; Li et al., 2017). Among the top five models generated by iterative simulations, the first one was selected for further analysis based on model quality evaluation using the VADAR version1.8 program (http://vadar.wishartlab.com) (Willard et al., 2003) and the ProSa-web program (https://prosa.services.came.sbg.ac.at/prosa.php)(Wiederstein and Sippl, 2007). Structures of point-mutated S-RNases were generated by Mutagenesis in PyMol. Electrostatic potential analysis of PhS_3_-RNase structure and its point-mutated ones were performed using plug-in APBS tools in PyMol as previously described (Baker et al., 2001; Li et al., 2017).

### Quantitative (q) RT-PCR analysis

Total RNAs were separately isolated from pistils derived from *PhS_3_S_3L_/PhS_3_-RNase-, /MI-, /MII-, /MK*-, /*MT*-, /*MIII*- and /*MI/II/III*-(*FLAG*) using TRIzol reagent (Ambion) according to the manufacturer’s instructions. cDNA was subsequently synthesized using TransScript-uni one-Step gDNA removal and cDNA synthesis supermix (Transgen, AU311-02). qRT-PCR reaction mixes were prepared according to manufacturer’s guidelines of ChamQ™ Universal SYBR qPCR Master Mix (Vazyme, Q711-02/03). Relevant primer sequences are listed in Supplemental Table 3. qRT-PCR assays were performed by CFX96™ Real-Time System (Bio-Rad). *P. hybrida 18S rRNA* gene trascripts were used as an internal control. The data were analyzed with the method of Livak (2^-ΔΔCt^) (Livak and Schmittgen, 2001).

### Mass spectrometry analysis for ubiquitination sites

The wild type self-incompatible *PhS_3_S_3L_* plants were self-pollinated or cross-pollinated with the pollen from the transgenic self-compatible plants *PhS_3_S_3L_/PhS_3L_SLF1*. Then 25 pollinated pistils were collected after 2, 6, 12 and 24 h, respectively. The pollinated pistils of the four time points were then mixed up, minced with liquid nitrogen and lysed in lysis buffer containing 7 M urea, 2 M thiourea and 0.1% CHAPS, followed by 5-minute ultrasonication on ice. Samples of unpollinated pistils were prepared as controls using the same strategy. The lysate was centrifuged at 14,000 g for 10 min at 4°C and the supernatant was transferred to a clean tube. Extracts from each sample were reduced with 10 mM DTT for 1 h at 56°C, and subsequently alkylated with sufficient iodoacetamide for 1 h at room temperature in the dark (Udeshi et al., 2013b). The supernatant from each sample containing precisely 10 mg of protein was digested with Trypsin Gold (Promega) at 1:50 enzyme-to-substrate ratio. After 16 h of digestion at 37°C, peptides were desalted with C18 cartridge to remove the high urea, and desalted peptides were dried by vacuum centrifugation (Udeshi et al., 2013b). The lyophilized peptides were resuspended with MOPS IAP buffer (50 mM MOPS, 10 mM KH_2_PO_4_ and 50 mM NaCl) adjusting to pH 7.0 with 1 M Tris, and centrifuged for 5 min at 12,000 g. Supernatants were mixed with anti-Ubiquitin Remnant Motif (K-ε-GG) beads (CST #5562, Cell Signaling Technology) for 2.5 h at 4°C, and then centrifuged for 30 s at 3,000 g at 4°C. Beads were washed in MOPS IAP buffer, then in water, prior to elution of the peptides with 0.15% TFA (Udeshi et al., 2013a). Then the peptides were desalted using peptide desalting spin columns (Thermo Fisher, 89852) before LC-MS/MS analysis on the Orbitrap Fusion mass spectrometer (Thermo Fisher). The resulting spectra from each fraction were searched separately against S-RNase amino acid sequences by the Maxquant search engines. Precursor quantification based on intensity was used for label-free quantification.

### S-RNase activity assays

The coding sequence of *PhS_3_-RNase* (without signal peptide) as well as its six mutant forms described above were separately cloned into *pCold-TF* vector (Takara). Relevant primer sequences are listed in Supplemental Table 3. Trigger Factor (TF) is a 48 kDa soluble tag located at the N-terminus of His. The His-tagged fusion proteins were respectively expressed in *Escherichia coli Trans* BL21 (DE3) plysS (Transgen) at 16°C for 24 h at 180 rpm, and then immobilized on Ni Sepharose 6 Fast Flow beads (GE Healthcare, 10249123) according to the manufacturer’s instructions. The beads were subsequently washed with wash buffer (25 mM pH 8.0 Tris-HCl, 150 mM NaCl, 15 mM imidazole), followed by an elution using buffer containing 25 mM pH 8.0 Tris-HCl, 150 mM NaCl, 250 mM imidazole. Protein concentration was determined by Bradford protein assay. The purified recombinant proteins were separately added into the tubes containing lyophilized fluorescent substrate according to the manufacture’s instructions of RNase Alert Lab Test Kit (Ambion). Each samples were then pipetted into a 96-well plate and incubated in Synergy 2 (Biotech) at 37°C. The relative fluorescence units (RFU) was monitored at 5 min internals for 2 h with 490 nm/520 nm excitation/emission wavelengths.

### Ubiquitination assay and immunoblotting

The SCF^SLF-FLAG^ complex attached to anti-FLAG M2 affinity gel (Sigma-Aldrich) serving as E3 ubiquitin ligase was purified from transgenic pollen tubes of *PhS_3_S_3L_/PhS_3L_SLF1-FLAG* as described (Li et al., 2017), with that from wild type *PhS_3_S_3L_* as a negative control. PhS_3_-RNase was purified from pistils of *PhS_3_S_3_* through Fast Protein Liquid Chromatography (FPLC) as described (Entani et al., 2014; Li et al., 2017), and recombinant His-PhS_3_-RNase and its six mutant forms described above was separately used as a substrate and added into mixture containing E1, E2, E3 on anti-FLAG gel, biotinylated ubiquitin and ATP for ubiquitination reaction (Li et al., 2017). After incubated at 37°C for 6 h, reaction was quenched by an addition of 2 × Non-reducing gel loading buffer (ChemCruz, B1919) and centrifuged at 6,000 g for 30 s. Supernatants were separated by 12% SDS-PAGE, transferred to PVDF (Millipore) and probed by primary antibodies including mouse monoclonal anti-PhS_3_-RNase, anti-ubiquitin (Abgent), and anti-His (Sigma) antibodies at a 1:2000 dilution, respectively. The PVDF membranes were washed with TBS-T buffer, followed an incubation with secondary antibody horseradish peroxidase (HRP)-conjugated goat anti-mouse IgG (Sigma) at a 1:10,000 dilution. Then the PVDF membrane were washed again with TBS-T buffer, and then the HRP signal were detected by Image Quant LAS4000 or Tanon5800 after incubated with Immoblion Western Chemiluminescent HRP substrate (Millipore). Image J was used to quantify the ubiquitinated recombinant His-PhS_3_-RNase (or its mutant forms) fusion proteins. K48-or K63-linkage Specific Polyubiquitin Rabbit mAb (Cell Signaling) at a 1:1000 dilution and the corresponding secondary antibody horseradish peroxidase (HRP)-conjugated goat anti-rabbit IgG (Sigma) at a 1:10,000 dilution were used to analyze the linkage type of PhS_3_-RNase ubiquitination.

### Subcellular fractionation and immunoblotting

Mature pollen grains of *PhS_V_S_V_* were suspended and cultured in liquid pollen germination medium (LPGM) as described (Liu et al., 2014) for 2-3 h, and then collected by centrifugation at 1,000 g for 2 min with the supernatant discarded. Stylar lysates extracted from transgenic pistils containing PhS_3_-RNase-FLAG or its six mutant forms by fresh LPGM were separately used to further incubate the collected pollen tubes. After cultured for 1 h, the treated pollen tubes were harvested for fractionation and equal amount of protein samples derived from each step of centrifugation were applied to immunoblotting as described (Liu et al., 2014).

### Cell-free degradation assays

Germinated pollen tubes of *PhS_V_S_V_* were collected and ground into fine powder in liquid nitrogen. Then total proteins were extracted on ice using cell-free degradation buffer containing 25 mM Tris-HCl (pH 7.5), 10 mM NaCl, 10 mM MgCl_2_, 1 mM DTT, 10 mM ATP and 1 mM PMSF, followed with a centrifugation at 12,000 g at 4°C for 15 min. The supernatants were subsequently quantified by Bradford method and equal amount of total proteins were applied to react with recombinant SUMO-His-PhS_3_-RNase or its mutant forms in the presence or the absence of 40 μM MG132. During incubation at 30°C, equal amount of samples were taken out at indicated time points to detect SUMO-His-tagged protein abundance through immunoblotting. Image J was used to quantify the results. The SUMO-His-tagged fusion proteins were generated as follows. The coding sequence of *PhS_3_-RNase* (without signal peptide) as well as its six mutant forms described above were separately cloned into engineered *pET-30a* (Novagen) containing N-terminal SUMO tag to produce SUMO-His-tagged proteins. Relevant primer sequences are listed in Supplemental Table 3. The fusion proteins were respectively expressed in *Escherichia coli* Trans BL21 (DE3) plysS (Transgen) at 16°C for 24 h, and then immobilized on Ni Sepharose 6 Fast Flow beads (GE Healthcare, 10249123). The beads were subsequently washed and eluted by buffers as described above.

### MBP pull-down assays

The coding sequences of *PhS_3_-RNase* (without signal peptide) and its six mutant forms were separately cloned into *pMAl-c2x* (Novagen) to generate MBP-tagged fusion proteins. The full length of *PhS_3L_SLF1* was cloned into engineered *pET-30a* (Novagen) described above to produce SUMO-His-PhS_3L_SLF1. Relevant primer sequences are listed in Supplemental Table 3. All the recombinant proteins were induced overnight at 16°C at 180 rpm, with MBP-tagged proteins expressed in *E. coli* Trans BL21 (DE3) plysS cells described above and SUMO-His-PhS_3L_SLF1 in *E. coli* Trans BL21 (DE3) (Transgen). Cells were subsequently collected and resuspended using binding buffer [20 mM Tris-HCl (pH 8.0), 200 mM NaCl, 1 mM DTT and 1 mM EDTA (pH 8.0)] for ultrasonication on ice. Then the lysates containing SUMO-His-PhS_3L_SLF1 were divided into seven equal aliquots and rotarily incubated with the same amount of lysates containing MBP or MBP-tagged fusion protein (PhS_3_-RNase or its six mutant forms), respectively for 2 h at 4°C. The mixed lysates were subsequently immobilized on Dextrin Sepharose High Performance (GE heathcare, 10284602) following the manufacturer’s instructions. Then the beads were washed with binding buffer and eluted by binding buffer supplemented with 10 mM maltose. The elutes were separated by 12% SDS-PAGE and subjected to immunoblotting using anti-MBP (NEB) and anti-His (Sigma) antibodies.

### Bimolecular fluorescence complementation (BiFC) assays

The coding sequences of *PhS_3_-RNase* (without signal peptide) and its six mutant forms were separately amplified and inserted into *pSY-735-35S-cYFP-HA* and the full-length cDNA of *PhS_3L_SLF1* was cloned into *pSY-736-35S-nYFP-EE* as described (Li et al., 2018). Relevant primer sequences are demonstrated in Supplemental Table 3. Different types of constructs (e.g., *nYFP-PhS_3L_SLF1* and *cYFP-PhS_3_-RNase*), together with *p19* silencing plasmid, were cotransfected into tobacco leaf epidermal cells by *Agrobacterium* (GV3101)-mediated infiltration to generate fusion proteins (e.g., nYFP-PhS_3L_SLF1 and cYFP-PhS_3_-RNase) for their interaction test. After cultured another 48 h in the dark, a portion of the injected leaf was cut off and subjected to confocal microscope (Zeiss LSM710) to capture the YFP signal.

### Split firefly luciferase complementation (SFLC) assays

The coding sequence of *PhS_3L_SLF1* (or *PhS_3_SLF1*) and *PhS_3_-RNase* (or its mutant forms) were cloned into *pCAMBIA1300-35S-HA-nLUC-RBS* and *pCAMBIA1300-35S-cLUC-RBS* vectors, respectively, as described (Liu et al., 2018). Relevant primer are listed in Supplemental Table 3. Then different construct combinations (e.g., *PhS_3L_SLF1-nLUC* and *cLUC-PhS_3_-RNase*) together with *p19* silencing plasmid were cotransfected into tobacco leaf epidermal cells via GV3101 described above. After 48 h in the dark, 1 mM luciferin was sprayed on the injected leaves with a 5-minute dark incubation. Then capture the LUC signal using a cooled CCD imaging system (Berthold, LB985).

### Aniline blue staining of pollen tubes within pistils

After self-pollination of wild-type plant *PhS_3_S_3L_* and transgenic lines containing *PhS_3L_SLF1-FLAG*, the pollinated pistils were collected and chemically fixed in ethanol: glacial acetic acid (3:1) solution for 24 h at 4°C. Then treat the pistils sequentially with 8 N sodium hydroxide, water and aniline blue solution to stain the pollen tubes within pistils for observation as described (Liu et al., 2014).

### Accession numbers

Sequence data presented in Supplemental Figure 3 can be found in the GenBank data library under the following accession numbers: *Petunia hybrida* S_3_-RNase (U07363), PhS_3L_-RNase (AJ271065), PhS_V_-RNase (AJ271062), *Pyrus bretschneideri* S_7_-RNase (XM_009350009), *Petunia inflata* S_2_-RNase (AAG21384), PiS_3_-RNase (AAA33727), PiS_k1_-RNase (BAE73275), PiS_1_-RNase (AAA33726), *Solanum lycopersicum* S_3_-RNase (XP_004229063), *Nicotiana alata* S_A2_-RNase (AAA87045), PhS_X_-RNase (AAA33729), *Petunia axillaris* S_C2_-RNase (AAN76454), *Solanum tuberosum* S_2_-RNase (Q01796), PhS_B2_-RNase (BAA76514), *Solanum neorickii* S-RNase (BAC00940), *Solanum habrochaites* S_2_-RNase (AIG62995), *Solanum chilense* S_1_-RNase (BAC00934), *Solanum chacoense* S_11_-RNase (AAA50306), ScS_14_-RNase (AAF36980), NaS7-RNase (Q40381), PaS_C1_-RNase (AAN76453), PhS_11_-RNase (BAJ24848), PhS_7_-RNase (BAJ24847), PaS_1_-RNase (AAK15435), ShS_4_-RNase (AIG62997), PhS_1_-RNase (AAA60465), PhS_B1_-RNase (BAA76513), *Solanum peruvianum* S_22_-RNase (BAC00930), SpS_12_-RNase (AAA77040), ShS_1_-RNase (AIG62994), PhS_0_-RNase (ACT35737), ScS_12_-RNase (AAD56217), NaS_6_-RNase (AAB40028), SpS_3_-RNase (CAA53666), SpS_11_-RNase (AAA77039), NaS_2_-RNase (P04007), and PaS_13_-RNase (AAK15436).

## ACKNOWLEDGMENTS

This work was supported by the Strategic Priority Research Program of the Chinese Academy of Sciences (XDB27010302) and the National Natural Science Foundation of China (32030007).

## AUTHOR CONTRIBUTIONS

Y.X conceived and designed the project. H.Z. and Y.S. performed the experiments. J.L. and Y.Z. conducted functional analyses of *PhS_3L_SLF1.* H.H. assisted transgenic plant construction. Q.L. and Y.E.Z. provided technical support. H.Z. and Y.X. analyzed data and wrote the manuscript. All authors commented on the article.

## Supplemental Data

**Supplemental Figure 1.** Identification of six ubiquitinated residues of PhS_3_-RNase by LC-MS/MS analysis of cross-pollinated pistils.

**Supplemental Figure 2.** Two ubiquitinated residues of PhS_3_-RNase identified by LC-MS/MS analysis of self-pollinated and unpollinated pistils.

**Supplemental Figure 3.** Locations of six ubiquitinated residues of PhS_3_-RNase in Solanaceous S-RNases.

**Supplemental Figure 4.** PhS_3_-RNase with the mutated R I displays largely unaltered biochemical and physical properties.

**Supplemental Figure 5.** Physical interactions between S_3_R/M and PhS_3_SLF1.

**Supplemental Figure 6.** Predicted 3D structures and surface electrostatic potentials of PhS_3_-RNases with mutated ubiquitinated residues.

**Supplemental Figure 7.** *S_3_R* and *S_3_R (M)* transgenes identification by PCR analysis.

**Supplemental Figure 8.** Identification of *FLAG*-tagged *S_3_R* and *S_3_R (M)* transgenes by PCR analysis.

**Supplemental Figure 9.** Detection of *FLAG-tagged S_3_R* and *S_3_R (M)* transgenes expression by immunoblots.

**Supplemental Figure 10.** PhS_3_-RNase with the mutated R I slightly reduces cross seed sets.

**Supplemental Figure 11.** Decreased ubiquitination amount of MI, II and III mediated by SCF^PhS_3L_SLF1^.

**Supplemental Figure 12.** Reduced seed set per capsule from T_0_ transgenic lines with mutated lysine or threonine within the six ubiquitinated residues of PhS_3_-RNase.

**Supplemental Figure 13.** MIII largely maintains the biochemical and physical properties of PhS_3_-RNase.

**Supplemental Figure 14.** PhS_3_-RNase with mutated R III markedly reduces cross seed sets.

**Supplemental Figure 15.** Largely unaltered physicochemical properties of PhS_3_-RNase with mutated R I, II and III.

**Supplemental Table 1.** Seed sets of *S_3_S_3L_/S_3_R* and *S_3_S_3L_/S_3_R (M)* T_0_ transgenic plants.

**Supplemental Table 2.** Seed sets of *S_3_S_3L_/S_3_R-FLAG* and *S_3_S_3L_/S_3_R (M)-FLAG* T_0_ transgenic plants.

**Supplemental Table 3.** List of primer sequences.

